# Sleep and Motor Control by a Basal Ganglia Circuit

**DOI:** 10.1101/405324

**Authors:** Danqian Liu, Chenyan Ma, Weitong Zheng, Yuanyuan Yao, Yang Dan

## Abstract

From invertebrates to humans, a defining feature of sleep is behavioral immobility(Campbell and Tobler, 1984; Hendricks et al., 2000; Shaw et al., 2000). In mammals, diminished electromyographic (EMG) activity is a major criterion for both rapid eye movement (REM) and non-REM (NREM) sleep. However, the relationship between sleep and motor control at the neuronal level remains poorly understood. Here we show that regions of the basal ganglia long known to be essential for motor suppression also play a key role in sleep generation. Optogenetic or chemogenetic activation of GABAergic neurons in mouse substantia nigra pars reticulata (SNr) strongly increased both REM and NREM sleep, whereas their inactivation suppressed sleep and increased wakefulness. Analysis of natural home-cage behavior showed that mice transition sequentially through several behavioral states: locomotion, non-locomotor movement, quiet wakefulness, and sleep. Activation/inactivation of SNr neurons promoted/suppressed sleep by biasing the direction of progression through the natural behavioral sequence. Virus-mediated circuit tracing showed that SNr GABAergic neurons project to multiple wake-promoting monoaminergic cell groups in addition to the thalamus and mesencephalic locomotor region, and activating each projection promoted sleep. Within the thalamus, direct optogenetic inactivation of glutamatergic neurons is sufficient to enhance sleep, but the effect is largely restricted to the regions receiving SNr projection. Furthermore, a major source of excitatory inputs to the SNr is the subthalamic nucleus (STN), and activation of neurotensin-expressing glutamatergic neurons in the STN also promoted sleep. Together, these results demonstrate a key role of the STN-SNr basal ganglia pathway in sleep generation and reveal a novel circuit mechanism linking sleep and motor control.

## Main

Sleep is characterized by a marked reduction of somatic motor activity(Campbell and Tobler, 1984; Hendricks et al., 2000; Shaw et al., 2000). Since the SNr GABAergic neurons play an important role in movement suppression(Gerfen and Surmeier, 2011; Hikosaka and Wurtz, 1985; Kravitz et al., 2010), we wondered whether they are also involved in sleep generation. A Cre-inducible adeno-associated virus (AAV) expressing channelrhodopsin 2 (ChR2) fused with enhanced yellow fluorescent protein (eYFP) was injected bilaterally into the SNr of *Slc32a1*^*Cre*^ mice to target the neurons expressing the vesicular GABA transporter (VGAT) (Fig. 1a, Supplementary Fig.1a). Laser stimulation (constant light, 2 min per trial) was applied randomly every 7-15 min in freely moving mice in their home cage, and wake, REM, and NREM sleep were classified based on electroencephalogram (EEG) and electromyogram (EMG) recordings. Activation of the SNr GABAergic neurons significantly increased both NREM (*P* < 0.0001, bootstrap) and REM (*P* < 0.0001) sleep and decreased wakefulness (*P* < 0.0001, Fig. 1b) by enhancing wake→NREM and NREM→REM transitions (Supplementary Fig.2a). In contrast, optogenetic inactivation of these neurons through a light-activated chloride channel (iC++)(Berndt et al., 2016) markedly increased wakefulness (*P* < 0.0001) and decreased sleep (Fig. 2c, Supplementary Fig.1b) by enhancing both NREM→wake and REM→wake transitions (Supplementary Fig.2b). In control mice expressing eYFP alone, laser stimulation had no effect (Supplementary Fig.2c), and the effects of laser stimulation were significantly different between the ChR2 and control mice (*P* < 0.0001, bootstrap) and between iC++ and control mice (*P* < 0.0001).

**Figure 1.**
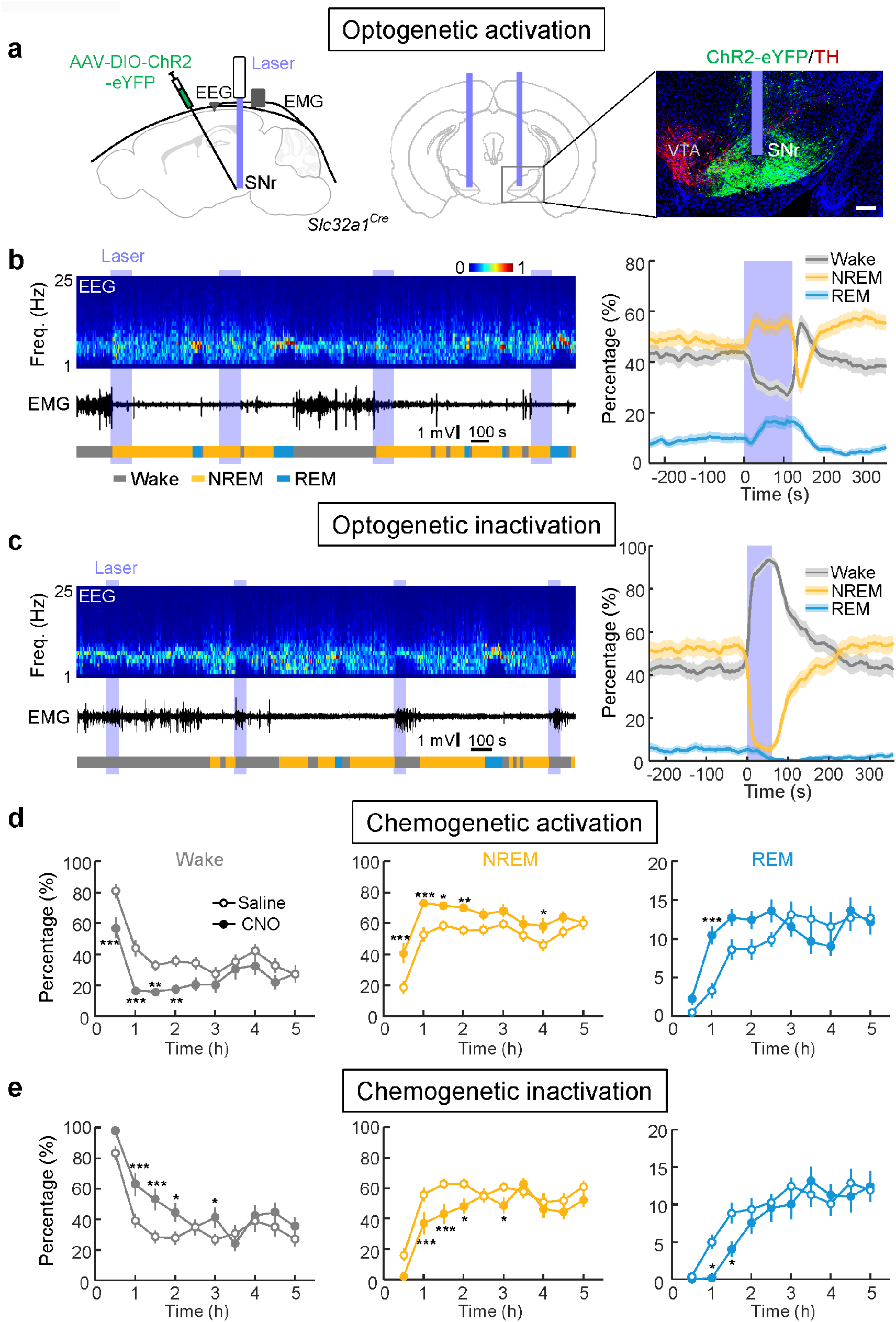
Activation and inactivation of SNr GABAergic neurons enhances and suppresses sleep, respectively. **a,** Left, schematic of ChR2-mediated activation of SNr GABAergic neurons. Right, fluorescence image of a region (black box in coronal diagram) in a *Slc32a1*^*Cre*^ mouse showing ChR2-eYFP expression (green) in SNr and tyrosine hydroxylase (TH) labelling (red) in the ventral tegmental area (VTA). Scale bar, 200 μm. Brain outline adapted from Allen Mouse Brain Atlas (available from: http://brain-map.org/api/index.html). **b,** Left, an example optogenetic activation experiment. Shown are EEG power spectrogram, EMG trace and brain states (color coded). Blue stripe, laser stimulation period (constant light, 120 s). Freq., frequency. Right, percentage of time in NREM, REM, or wake state before, during, and after laser stimulation, averaged from 9 mice. Shading, 95% confidence intervals (CI). **c,** Left, example optogenetic inactivation experiment. Blue stripe, laser stimulation period (constant light, 60 s). Right, percentage of time in NREM, REM, or wake state before, during, and after laser stimulation, averaged from 6 mice (NREM, *P* < 0.0001; REM, *P* = 0.01; wakefulness, *P* < 0.0001). **d,** Effect of chemogenetic activation of SNr. Shown is the percentage of time in each brain state following saline or CNO injection in mice expressing hM3Dq in the SNr (n = 6 mice). Horizontal axis, time after CNO/saline injection. Error bar, ±SEM. \**P* < 0.05; ** *P* < 0.01; *** *P* < 0.001 (two-way ANOVA with Bonferroni correction). **e,** Similar to **d**, for chemogenetic inactivation (n = 6 mice).

**Figure 2.**
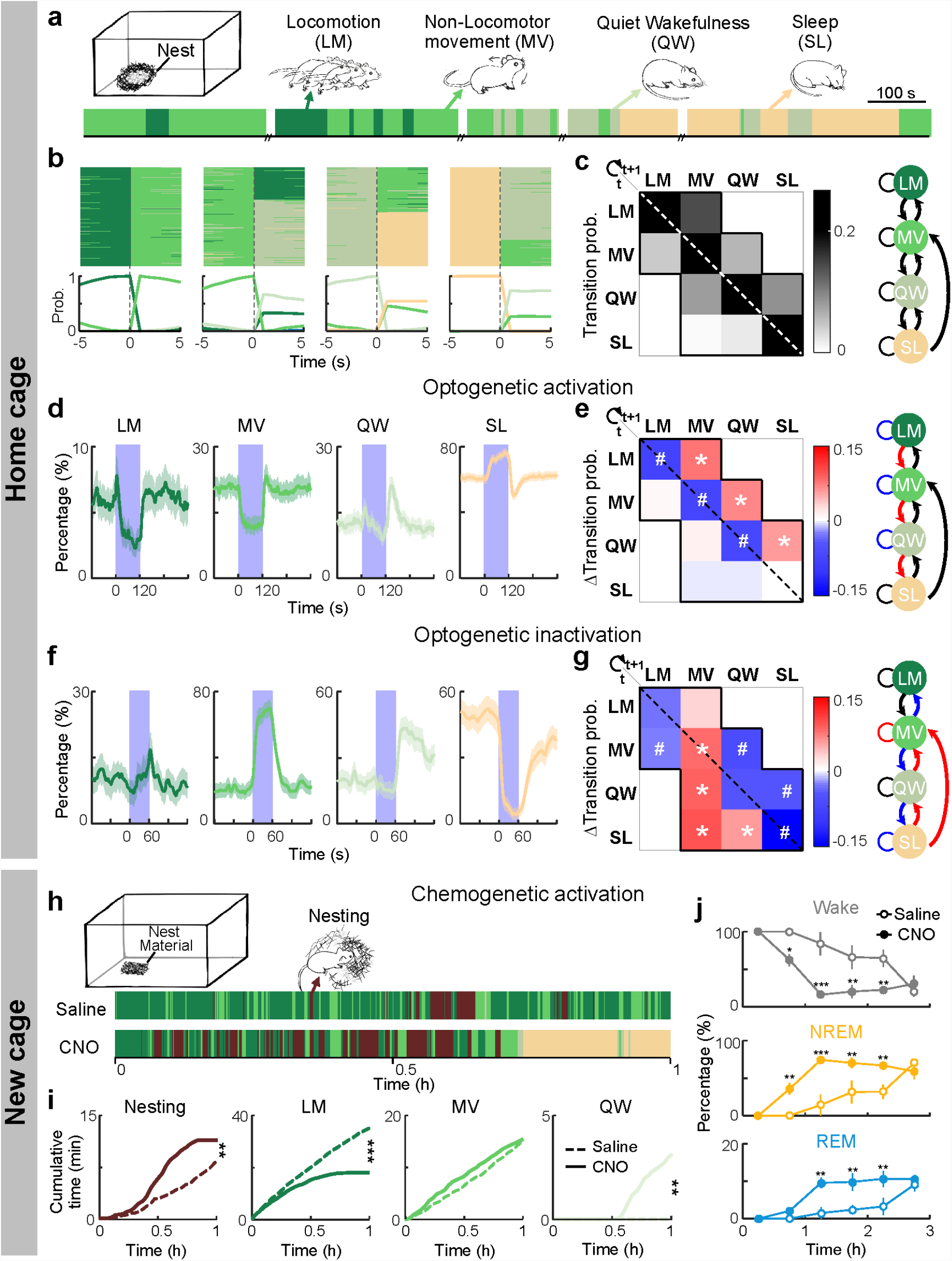
Effects of SNr GABAergic neuron activity on mouse motor behavior in home cage and new cage. **a,** Natural home-cage behaviors (color coded) in an example recording session. **b,** Top, all transitions from each behavior (LM, MV, QW or SL) recorded in 16 mice aligned to time of transition (t = 0). Bottom, probability of each behavior before and after transition. **c,** Left, transition probability between each pair of behaviors within each 5 s. Diagonal pixels indicate maintenance of each behavior. Right, summary of natural transitions. **d,** Percentage of time in each behavior before, during and after optogenetic activation of SNr neurons, averaged from 8 mice. The effect of laser was significant for LM, MV and SL (*P* < 0.0001, bootstrap). **e,** Left, changes in transition probabilities induced by SNr activation (difference between the 120-s period before and 120 s during laser stimulation). *, significantly increased; #, significantly decreased (*P* < 0.001, bootstrap). Right, summary of transition probabilities that are significantly increased (red), decreased (blue) or unchanged (black). **f-g,** Similar to **d-e**, for SNr inactivation (laser stimulation, 60 s; n = 4 mice). The effect of laser was significant for MV, QW and SL in **f** (*P* < 0.0001, bootstrap). **h,** Behaviors of a mouse expressing hM3Dq in SNr GABAergic neurons in a new cage (containing nest material) in example sessions after saline and CNO injections. Mice were transferred into a new cage 15 min after injection (t = 0, time of cage transfer). **i,** Cumulative time spent in each behavior after entering the new cage following CNO or saline injection. ** *P* < 0.01, *** *P* < 0.001, two-sample Kolmogorov-Smirnov test. n = 6 mice. **j,** Effect of SNr chemogenetic activation on wakefulness, NREM, and REM sleep in the new cage. Error bar, ±SEM. * *P* < 0.05, ** *P* < 0.01 (two-way ANOVA with Bonferroni correction).

We next tested the effects of chemogenetic manipulations of SNr GABAergic neurons using Designer Receptors Exclusively Activated by Designer Drug (DREADDs). In mice expressing the excitatory DREADD hM3Dq (Supplementary Fig.1c), clozapine-N-oxide (CNO) injection significantly increased NREM and REM sleep and decreased wakefulness (Fig. 1d). In contrast, in mice expressing hM4Di (inhibitory DREADD) in their SNr GABAergic neurons (Supplementary Fig.1d), CNO injection had the opposite effects (Fig. 1e). In control mice expressing mCherry only, CNO injection had no significant effect (Supplementary Fig.2d). Together, the bidirectional optogenetic and chemogenetic manipulation experiments demonstrate a crucial role of the SNr in sleep generation.

To understand how the SNr GABAergic neurons participate in both sleep generation and motor suppression(Gerfen and Surmeier, 2011; Hikosaka and Wurtz, 1985; Kravitz et al., 2010), we first analyzed the relationship between sleep and various motor behaviors of the mice in their home cage. Based on EEG, EMG, and video recordings, the home-cage behaviors were classified into locomotion (LM), non-locomotor movement (MV, including eating, grooming, and postural adjustments), quiet wakefulness (QW), and sleep (SL, Fig. 2a), ranked by decreasing levels of both motor activity and EEG activation(Crochet and Petersen, 2006; McGinley et al., 2015; Niell and Stryker, 2010; Polack et al., 2013; Poulet et al., 2012). Interestingly, almost all the behavioral transitions were between adjacent states, with a single exception of direct SL→MV transition (Fig. 2b, c).

We then analyzed the effects of SNr optogenetic manipulations. Along with the increase in SL, GABAergic neuron activation caused strong decreases in both LM and MV (Fig. 2d), consistent with their known function in movement suppression(Gerfen and Surmeier, 2011; Hikosaka and Wurtz, 1985; Kravitz et al., 2010). However, these effects were not caused by direct LM→SL or MV→SL transitions, which were not observed during natural behaviors; instead, laser stimulation significantly enhanced LM→MV, MV→QW, and QW→SL transitions (Fig. 2e, Supplementary Fig.3a, c, d), all of which occur naturally between adjacent states (Fig. 2b, c). Notably, no increase was observed in any transition in the opposite direction. Optogenetic inactivation of SNr GABAergic neurons, in contrast, significantly increased MV while suppressing SL, by decreasing MV→QW and QW→SL transitions and increasing SL→QW, SL→MV, and QW→MV transitions (Fig. 2f, g, Supplementary Fig.3b). Thus, without inducing artificial behavioral transitions, SNr activation/inactivation strongly biased the natural transitions in the direction of decreasing/increasing motor activity and EEG activation.

Transferring a mouse from the familiar home cage into a new cage is known to increase arousal and decrease sleep(Pawlyk et al., 2008; Suzuki et al., 2013). In addition to exploring the novel environment, the mouse typically builds a new nest before sleep(Eban-Rothschild et al., 2016). In mice expressing hM3Dq in their SNr GABAergic neurons, CNO injection caused a strong reduction of exploratory locomotion but an increase in nest-building activity during the first hour in the new cage (Fig. 2h, i). As a result, the nest quality was significantly higher than the control (Supplementary Fig.4a). This is similar to the effect of chemogenetic inactivation of dopaminergic neurons in the ventral tegmental area (VTA)(Eban-Rothschild et al., 2016), suggesting that the effect of SNr activation may be partially mediated by its inhibition of the dopaminergic neurons(Ogawa et al., 2014). Following the flurry of nest-building activity within the first hour, the hM3Dq-expressing mice injected with CNO spent ∼70% and ∼10% of the time in NREM and REM sleep, respectively, much higher than the control mice not expressing hM3Dq and/or injected with vehicle without CNO (Fig. 2j, Supplementary Fig.4b, c). Thus, instead of suppressing all physical movement indiscriminately, SNr activation in an unfamiliar environment causes a shift from exploration to a motor behavior naturally preceding and conducive to sleep.

Anterograde labeling of SNr neuron axons showed that in addition to several brain regions involved in motor control, including the thalamus, pedunculopontine nucleus (PPN), and superior colliculus (SC)(Grillner et al., 2005; Roseberry et al., 2016), the GABAergic neurons also project to multiple monoaminergic cell groups (Fig. 3a, b). Rabies virus (RV)-mediated retrograde transsynaptic tracing confirmed their direct innervation of VTA dopaminergic neurons, dorsal raphe (DR) serotonergic neurons, and locus coeruleus (LC) noradrenergic neurons(Ogawa et al., 2014; Weissbourd et al., 2014) (Supplementary Fig.5), all of which promote wakefulness and arousal(Carter et al., 2010; Eban-Rothschild et al., 2016; Monti and Jantos, 2008). Optogenetic activation of SNr GABAergic axons in each region promoted sleep (Fig. 3c-f), suggesting that the effect of SNr stimulation is mediated at least in part by inhibiting the wake-promoting monoaminergic populations.

**Figure 3.**
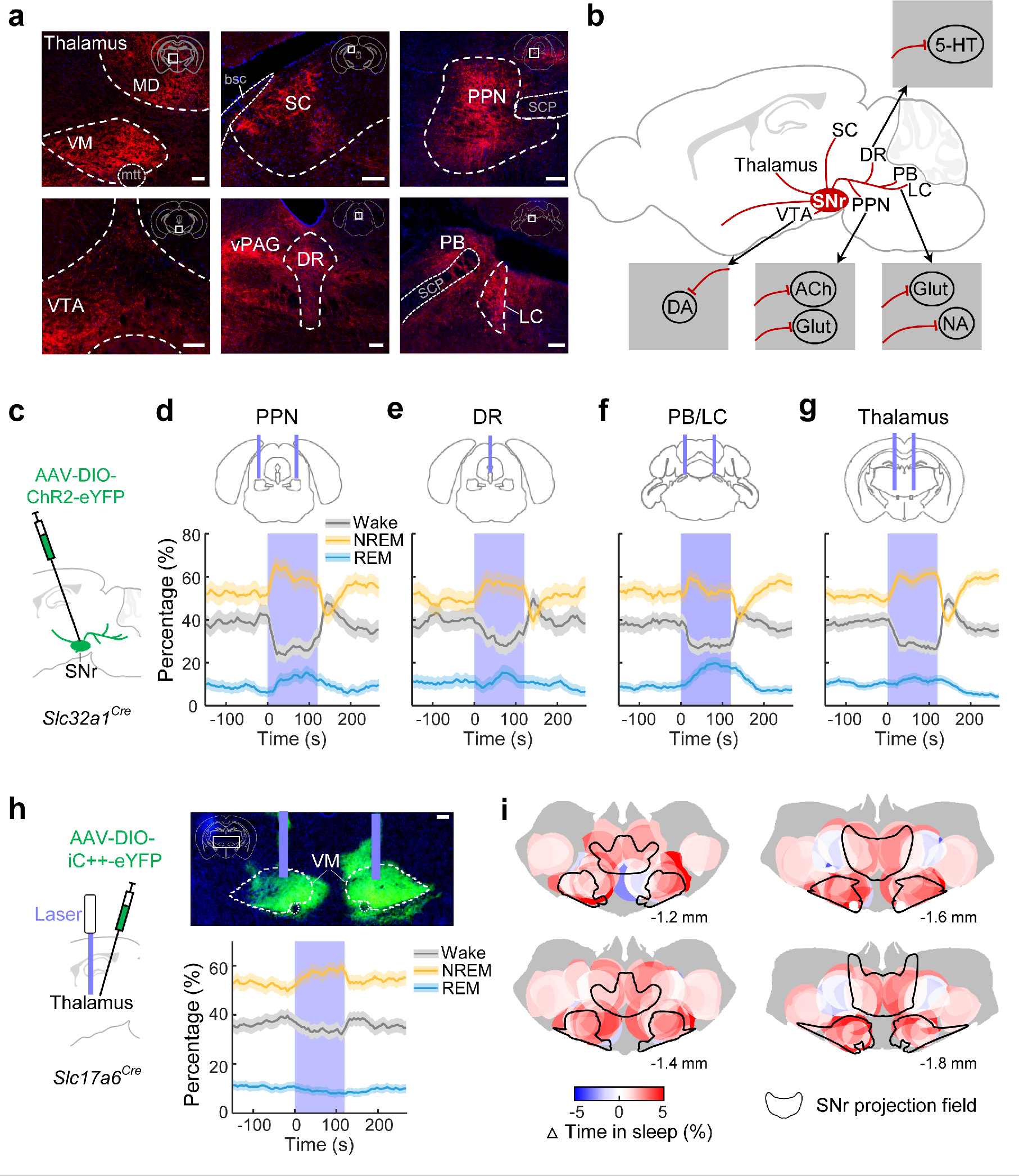
SNr GABAergic neurons promote sleep through multiple projections. **a,** Fluorescence images of several brain regions (white boxes in coronal diagrams) in a *Slc32a1*^*Cre*^ mouse injected with AAV-EF1α-DIO-mCherry into the SNr. Scale bars, 100 μm. VM/MD: ventral medial/medial dorsal thalamic nucleus; SC: superior colliculus; PPN: pedunculopontine nucleus; VTA: ventral tegmental area; DR: dorsal raphe; PB: parabrachial nucleus; LC: locus coeruleus; mtt: mammillothalamic tract; bsc: brachium superior colliculus; scp: superior cerebellar peduncles; vPAG: ventral periaqueductal gray. **b,** Summary of SNr GABAergic projections. Insets summarize neuronal populations receiving SNr monosynaptic innervation, shown by rabies-mediated transsynaptic tracing (Supplementary Fig.5). DA, dopaminergic; Glut, glutamatergic; ChAT, cholinergic; 5-HT, serotoninergic; NA, noradrenergic. **c,** Schematic of optogenetic stimulation of SNr axons. **d-g,** Percentage of time in NREM, REM, or wake state before, during, and after laser stimulation of SNr axons in PPN (**d**, n = 4 mice), DR (**e**, n = 4), PB/LC (**f**, n = 6), or thalamus (**g**, n = 7). **h,** Effect of optogenetic inactivation of glutamatergic neurons in a thalamic region receiving SNr input (n = 8). Scale bar, 200 μm. **i,** Effect of optogenetic inactivation on sleep (color coded) as a function of thalamic location (n = 26 mice). The effect at each location (combined change in REM and NREM sleep) was averaged from all the mice in which the given location exhibited iC++-eYFP expression and was within 500 μm from the optic fiber tip (see Methods and Supplementary Fig.8); gray indicates untested regions. Black outlines: areas with SNr projections, summed from 3 mice.

Notably, optogenetic activation of the SNr projection to the thalamus also enhanced NREM sleep (Fig. 3g), which could be mediated by the inhibition of thalamic neuron activity (Supplementary Fig.6). However, the thalamus-projecting SNr neurons also send axon collaterals to other wake-promoting brain regions (Supplementary Fig.7), and the effect of terminal stimulation in the thalamus could be mediated by antidromic activation of the SNr neurons and their axon collaterals in other brain areas. To test whether direct inactivation of thalamic glutamatergic neurons is sufficient to promote sleep, we injected Cre-inducible AAV expressing iC++ into the thalamus of *Slc17a6*^*Cre*^ mice, in a region that receives strong SNr projection (Fig. 3h). Laser stimulation induced a small but significant increase in NREM sleep (*P* < 0.001) and decrease in wakefulness (*P* < 0.001), indicating that direct suppression of thalamic spiking can contribute to sleep generation. Since the wake-promoting effect of thalamic activation varies across regions(Gent et al., 2018; Honjoh et al., 2018), we also measured the effect of optogenetic inactivation at a range of thalamic locations. The sleep-promoting effect was restricted to some midline and intralaminar thalamic nuclei believed to be part of the arousal system(Van der Werf et al., 2002) (Fig. 3i, Supplementary Fig.8), consistent with the previous finding based on optogenetic activation(Gent et al., 2018; Honjoh et al., 2018). Interestingly, the thalamic regions in which optogenetic manipulations are effective in brain state modulation correspond largely to the regions receiving strong SNr projection (Fig. 3i, black outline), suggesting that an important function of the SNr-thalamus projection is brain state regulation.

In principle, excitatory neurons upstream of the SNr could also promote sleep through their SNr projection. To screen for candidate sleep neurons along this pathway, we performed RV-mediated transsynaptic tracing from the GABAergic neurons. Fluorescence in situ hybridization (FISH) of *Slc17a6* revealed glutamatergic inputs from several brain regions, with the largest fraction originating from the subthalamic nucleus (STN)(Gerfen and Surmeier, 2011) (Fig. 4a-c, Supplementary Fig.9a). Within the STN, virtually all RV-labeled neurons expressed the gene encoding the peptide neurotensin (*Nts,* Fig. 4c), which labels STN neurons with a much higher specificity than *Slc17a6*. We thus targeted the STN neurons using *Nts*^*Cre*^ mice(Leinninger et al., 2011). ChR2-mediated activation of the STN neurons (50 Hz, 2 min/trial) caused a strong increase in NREM sleep and decrease in wakefulness (Fig. 4d, Supplementary Fig.9b, c). Similar changes were observed when laser stimulation was applied to the STN axons in the SNr (Fig. 4e), suggesting that the effect is largely mediated by the STN-SNr projection.

**Figure 4.**
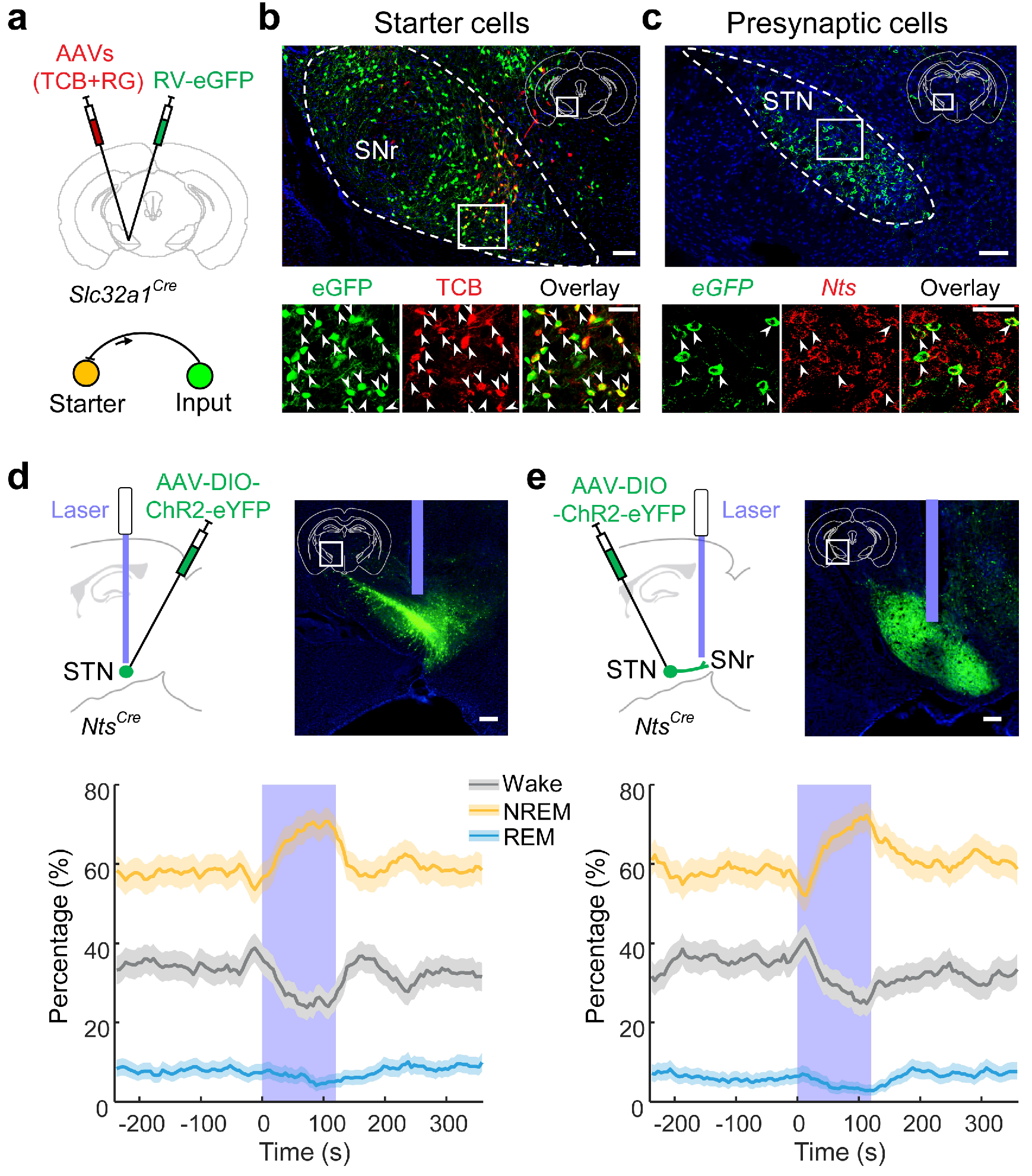
Glutamatergic neurons in STN innervate SNr GABAergic neurons and promote sleep. **a,** Schematic for rabies virus (RV)-mediated transsynaptic tracing from SNr GABAergic neurons. TCB: EnvA receptor fused with mCherry; RG: rabies glycoprotein. **b,** Top, fluorescence image of SNr in a *Slc32a1*^*Cre*^ mouse (white box in coronal diagram). Scale bar, 100 μm. Bottom, enlarged view of the region in white box showing starter cells (expressing both eGFP and mCherry, indicated by arrowheads). Scale bar, 50 μm. **c,** Top, RV-eGFP-labelled presynaptic neurons in STN. Scale bar, 100 μm. Bottom, enlarged view of the region in white box showing eGFP-labeled neurons expressing neurotensin (*Nts*). Arrowheads, overlap between *eGFP* and *Nts*. Scale bar, 50 μm. **d,** Percentage of time in NREM, REM, or wake state before, during, and after optogenetic activation of neurotensin-expressing STN neurons (n = 6 mice). Laser stimulation (50 Hz, 120 s) significantly increased NREM sleep (*P* < 0.0001, bootstrap) and decreased wakefulness (*P* < 0.0001) and REM sleep (*P* = 0.02). Shading, 95% CI. Scale bar, 200 μm. **e,** Percentage of time in NREM, REM, or wake state before, during, and after optogenetic activation of STN axons in SNr (n = 6 mice). Laser stimulation (50 Hz, 120 s) significantly increased NREM sleep (*P* < 0.0001, bootstrap) and decreased wakefulness (*P* = 0.0004) and REM sleep (*P* = 0.01). Shading, 95% CI. Scale bar, 200 μm.

## Discussion

Our finding on the role of SNr in sleep regulation (Fig. 1) is consistent with previous lesion studies in rats(Gerashchenko et al., 2006) and cats(Lai et al., 1999). Lesion of the globus pallidus also caused severe insomnia in rats(Qiu et al., 2010), which could be due to the loss of GABAergic neurons in the internal segment, a region with strong functional similarity to the SNr(Gerfen and Surmeier, 2011). Chemogenetic activation of adenosine A2A receptor-expressing neurons in the nucleus accumbens(Oishi et al., 2017) or dorsal striatum(Yuan et al., 2017), which can activate the STN and SNr neurons through the indirect pathway(Gerfen and Surmeier, 2011; Kravitz et al., 2010), has been shown to increase NREM sleep. However, unlike the effect of SNr neuron activation, which rises rapidly within ∼20 s of laser onset (Fig. 1b), sleep induction by optogenetic activation of the nucleus accumbens occurs gradually over ∼10 min of continuous laser stimulation(Oishi et al., 2017). A subset of GABAergic neurons in the zona incerta, a structure immediately adjacent to the STN, also promotes sleep(Liu et al., 2017). Similar to the SNr, the zona incerta projects to the thalamus and several wake-promoting monoaminergic cell groups(Halassa and Acsady, 2016; Liu et al., 2017), suggesting that the two GABAergic populations promote sleep through shared downstream pathways.

The LM→MV→QW→SL behavioral sequence of the mice observed in their home cage (Fig. 2a-c) bears a general resemblance to human behavior prior to sleep. Compared to QW and SL, LM and MV are associated with not only higher EMG activity, but also desynchronized EEG and stronger sensory responses indicative of heightened mental alertness(Crochet and Petersen, 2006; McGinley et al., 2015; Niell and Stryker, 2010; Polack et al., 2013; Poulet et al., 2012). These mental and physical changes may be coordinated by monoaminergic neurons, many of which regulate body movement as well as cortical activation(da Silva et al., 2018; Holstege and Kuypers, 1987; White et al., 1991). The PPN also controls both locomotion through its descending projection to the spinal cord and cortical activation through ascending projection to the midbrain and forebrain(Caggiano et al., 2018; Galtieri et al., 2017; Kroeger et al., 2017; Lee et al., 2014; Roseberry et al., 2016). The natural behavioral sequence leading to sleep could thus be mediated by a progressive inactivation of the monoaminergic and PPN neurons, resulting in both EEG synchronization and movement reduction. Indeed, the PPN, VTA, DR, and LC are all projection targets of the SNr (Fig. 3a, b), and these GABAergic projections contribute strongly to the sleep-promoting effect of SNr neuron activation (Fig. 3c-f).

Importantly, optogenetic activation of the SNr during wakefulness did not cause abrupt cessation of all motor activity, unlike that observed during cataplexy or that induced by activating glycinergic inputs from the pontine reticular nucleus to the thalamus(Giber et al., 2015). Instead, SNr activation suppressed or enhanced motor behaviors in a context-dependent manner: in the home cage it increased the LM→MV transition (Fig. 2e), whereas in a new cage it decreased exploration and increased nest building (Fig. 2h, i). In addition, SNr activation enhanced the QW→SL and NREM→REM transitions (Fig. 2e, Supplementary Figs.2a, S3a), which are mainly associated with EEG changes rather than movement reduction, indicating a role of the SNr in brain state regulation beyond simple motor suppression. Of course, the SNr GABAergic neurons are known to exhibit functional heterogeneity(Jin et al., 2014), and different neurons may be preferentially involved in motor command execution vs. brain state regulation(Dudman and Krakauer, 2016). The thalamus-projecting SNr neurons with axon collaterals to the PPN and wake-promoting monoaminergic regions (Supplementary Fig.7) seem to be well suited for regulating the global brain state through their divergent projections. Beyond the SNr, inhibitory neurons in the ventrolateral medulla can also suppress locomotor activity(Capelli et al., 2017) as well as promote REM sleep(Weber et al., 2015). Thus, the engagement of overlapping neuronal populations for motor suppression and sleep generation appears to be a common theme along the neural control pathway. Such a circuit design can ensure a strong coupling between sleep and immobility at the behavioral level, and breakdown of the shared control mechanism may underlie a variety of sleep-related movement disorders(Bargiotas and Bassetti, 2017).

## Methods

### Animals

All procedures were approved by the Animal Care and Use Committees of the University of California, Berkeley. Optogenetic manipulation experiments were performed in *Slc32a1*^*Cre*^ (Jackson stock no. 016962), *Slc17a6*^*Cre*^ (016963), and *Nts*^*Cre*^ (017525) mice. Rabies-mediated transsynaptic tracing experiments were performed in *Slc32a1*^*Cre*^, *Slc17a6*^*Cre*^, *Chat*^*Cre*^ (006410), *Sert*^*Cre*^ (014554), and *TH*^*Cre*^ (European Mouse Mutant Archive, EM00254) mice. Mice were housed in 12-hour (h) light-dark cycle (lights on at 07:00 am and off at 07:00 pm) with free access to food and water. Animals with implants were housed individually.

### Surgical procedures

Adult mice (6-12 weeks old) were anesthetized with 1.5-2% isoflurane and placed on a stereotaxic frame. Body temperature was kept stable throughout the procedure using a heating pad. Viruses were injected using a glass micropipette with a tip diameter of 15-20 μm, through a small skull opening (< 0.5 mm^2^), with Nanoject II (Drummond Scientific). For optogenetic manipulations, AAV2-EF1α-DIO-ChR2-eYFP, AAV2-EF1α-DIO-iC++-eYFP, or AAV2-EF1α-DIO-eYFP (∼5×10^12^ genome copies (gc)/mL, University of North Carolina Vector Core) was injected bilaterally into target regions (100-200 nl). For chemogenetic manipulations, AAV2-EF1α-DIO-hM3Dq-mCherry, AAV2-EF1α-DIO-hM4Di-mCherry, or AAV2-EF1α-DIO-mCherry (∼3**×**10^12^ gc/mL, University of North Carolina Vector Core) was injected bilaterally into the SNr (200-300 nl). For anterograde tracing, 150 nl of AAV2-EF1α-DIO-mCherry was injected unilaterally into the SNr.

Two to three weeks after virus injection, we implanted electroencephalogram (EEG) and electromyogram (EMG) electrodes for polysomnographic recordings. A reference screw was inserted into the skull on top of the right cerebellum. EEG recordings were made from two screws, one on top of hippocampus - anteroposterior (AP) -2.5 mm, mediolateral (ML) 2-2.5 mm - and one on top of the frontal cortex - AP 1.0 mm, ML 1.5 mm. Two EMG electrodes were inserted into the left and right neck muscles. For optogenetic experiments, optical fibers (0.2 mm diameter) were inserted bilaterally with the tip 0.2-0.5 mm above the viral injection sites (SNr, thalamus or STN). To stimulate axon projections of SNr GABAergic neurons, optical fibers were implanted bilaterally ∼0.2 mm above each region (except for DR at the midline, where a single fiber was implanted). Dental cement was applied to cover the exposed skull and secure the screws, electrodes and optical fibers.

Stereotaxic coordinates:

SNr: AP -3.0 mm, ML 1.4 mm, dorsoventral (DV) 4.6 mm.

Thalamus: AP -0.6 to -1.8 mm, ML 0.3 to 1.8 mm, DV 3.3 to 4.2 mm.

(VM: AP -1.5mm, ML 0.9 mm, DV 4.2 mm).

STN: AP -2.0 mm, ML 1.7 mm, DV 4.6 mm.

VTA: AP -2.8 mm, ML 0.3 mm, DV 4.5 mm.

PPN: AP -4.5 mm, ML 1.2 mm, DV 3.5 mm.

PB: AP -5.2 mm, ML 1.2 mm, DV 3.2 mm.

DR: AP -4.3, ML 0 mm, DV 3.2 mm.

LC: AP -5.3 mm, ML 0.9 mm, DV 3.5 mm.

### Rabies virus tracing

For retrograde transsynaptic tracing, AAV2-CAG-FLEx-TCB (∼5 × 10^12^ gc/mL) or AAV2-CAG-FLEx-TC^66T^ (∼5 × 10^12^ gc/mL, for VTA) and AAV2-CAG-FLEx-RG (∼5 × 10^12^ gc/mL, RG, rabies glycoprotein) were mixed at a 1:1 ratio and injected unilaterally into the target brain region (200-300 nl). Since VTA is close to the SNr, AAV-TC^66T^ was used instead of AAV-TCB for tracing from VTA to avoid local contamination(Miyamichi et al., 2013). Two to three weeks later, EnvA-pseudotyped, RG-deleted, and eGFP-expressing rabies virus (RV-eGFP, 300 nl at 1.0 × 10^9^ cfu/ml) was injected into the same location, and mice were sacrificed 5-7 days later. RV-eGFP was purchased from Gene Transfer Targeting and Therapeutics Core of Salk Institute, amplified in B7GG cells, pseudotyped with BHK-EnvA cells, and titered with HEK293-TVA cells.

For labelling the collaterals of thalamus-projecting SNr neurons, 300 nl AAV2-CAG-FLEx-TCB was injected into the SNr of *Slc32a1*^*Cre*^ mice. Three weeks later, 200 nl RV-eGFP was injected into the thalamus (VM), and mice were sacrificed 7 days later.

### Histology and immunohistochemistry

Mice were deeply anesthetized and transcardially perfused with 0.1 M phosphate-buffered saline (PBS) followed by 4% paraformaldehyde (PFA) in PBS. After dissection, brains were post-fixed overnight in 4% PFA and then dehydrated in 30% sucrose (in PBS) for over 1 day. Coronal sections were serially cut at 50 μm using a cryostat (Leica). Fluorescence images were taken using Nanozoomer 2.0 RS (Hamamatsu).

To determine the locations of virus expression in optogenetic and chemogenetic experiments in SNr and STN (Supplementary Figs.1, 9), coronal images were downsampled to 10 μm/pixel and aligned to templates of the Allen Mouse Brain Atlas (© 2015 Allen Institute for Brain Science. Allen Brain Atlas API. Available from: http://brain-map.org/api/index.html) using several anatomical landmarks. Viral expression areas were determined using ImageJ as those with green fluorescence signal above a threshold (20% of the maximal intensity). Area with obvious autofluorescence signals were excluded manually. All brain outline figures shown together with the histology images are adapted from Allen Mouse Brain Atlas.

To determine the effect of optogenetic inactivation of thalamic neurons as a function of location (Fig. 3i), iC++-eYFP expression within 500 μm from the optic fiber tip was mapped for each mouse. Since laser stimulation normally causes eYFP photobleaching, we sacrificed the mice immediately after the last optogenetic testing session and used the center of the photobleached area together with the optic fiber track to determine the location of the fiber tip. After mapping the viral expression area based on fluorescence, the photobleached area was manually filled. Signals outside the sphere (500 μm in radius, along AP, ML and DV axes) surrounding the fiber tip were excluded. The effect of optogenetic inactivation for each mouse was calculated as the difference in sleep (NREM + REM) time between the laser stimulation period and 0-300 s before laser stimulation. For each location in the thalamus we averaged the effect across all the mice in which iC++-eYFP was expressed at that location.

Immunohistochemistry (IHC) for GFP (chicken anti-GFP antibodies, Aves Labs, Cat#GFP-1020, 1:500) or TH (rabbit anti-TH antibodies, abcam, ab112, 1:500) was conducted as previously described(Chung et al., 2017). Fluorescence images were taken using a fluorescence microscope (Keyence, BZ-X710) and Nanozoomer 2.0 RS (Hamamatsu).

### Fluorescence in situ hybridization (FISH)

For rabies tracing experiments, we performed FISH to detect *Gad1/2* or *Slc17a6* mRNA. Two methods were used.

*RNAscope assays:* Brain sections were cut at 30 μm thickness using a cryostat, and double FISH was performed using RNAscope assays according to the manufacturer’s instructions (Advanced Cell Diagnostics). For RV tracing from GABAergic neurons in the SNr, we performed double FISH of *eGFP* and *Slc17a6 or Nts.* For RV tracing from dopaminergic neurons in the VTA, serotonergic neurons in the DR and glutamatergic neurons in the PPN, we performed double FISH of *Gad2* and *eGFP*.

*FISH-IHC:* For RV tracing from cholinergic neurons in the PPN, glutamatergic neurons in the PB and noradrenergic neurons in the LC, we performed FISH-IHC to detect *Gad1/2* mRNA and eGFP protein. FISH-IHC procedures were performed as previously described(Chung et al., 2017). In brief, sections were pretreated with 2% hydrogen peroxide and proteinase K (10 μg/ml) and then incubated with *Gad1* (forward: 5’-TGTGCCCAAACTGGTCCT -3’, reverse: 5’-TGGCCGATGATTCTGGTT -3’) or *Gad2* (forward: 5’-TCTTTTCTCCTGGTGGCG -3’, reverse: TTGAGAGGCGGCTCATTC - 3’) antisense riboprobes, for 16-20 h at 53-58 °C in a hybridization buffer. Sections were then washed with wash buffers and treated with RNase A (10 μg/ml). Immunochemistry staining for GFP was then performed as described above.

### Polysomnographic recordings

Behavioral experiments were carried out in home cages or new cages with nest materials placed in sound-attenuating boxes between 9:00 am and 7:00 pm. EEG and EMG electrodes were connected to flexible recording cables via a mini-connector. Recordings started after 20-30 min of habituation. The signals were recorded with a TDT RZ5 amplifier, filtered (0–300 Hz) and digitized at 1,500 Hz. Spectral analysis was carried out using fast Fourier transform (FFT), and brain states were classified into NREM, REM and wake states in 5 s epochs (wake: desynchronized EEG and high EMG activity, NREM: synchronized EEG with high-amplitude, low-frequency (1-4 Hz) activity and low EMG activity, REM: high power at theta frequencies (6-9 Hz) and low EMG activity). The classification was made using custom-written MATLAB software.

### Behavioral monitoring and classification

The experimenter analyzing the mouse behavior was blind to the experimental conditions. Mouse behavior was recorded using a camera placed on top of the cage, along with EEG and EMG recordings. Video frames during the wakeful state (based on polysomnographic recordings) were extracted for further behavioral annotation. We then performed second-by-second video analysis. In the home cage, waking behaviors were classified into three states: locomotion (LM), non-locomotor movements (MV, including grooming, eating and posture changes) and quiet wakefulness (QW, no movements). In the new cage with nest material, waking behaviors were classified into four states: LM, NL, QW and nesting (manipulating nest materials). The behavioral annotation was performed manually using a custom written graphical user interface programmed in MATLAB. Nest score was scored from 0 (poor) to 5 (good)(Eban-Rothschild et al., 2016), at 0.5 and 1 hour after the mouse enters a new cage.

### Optogenetic manipulation

Optic fiber was attached through an FC/PC adaptor to a 473-nm blue laser diode (Shanghai laser), and light pulses were generated using the TDT system. For SNr neurons, constant light (2 – 4 mW at fiber tip) lasting for 120 s (activation) or 60 s (inactivation) was used. For thalamic neurons, constant light (2 - 4 mW, 120 s) was used. For STN neurotensin-expressing neurons, high frequency laser (50 Hz, 10 ms per pulse, 1 mW, 120 s) was used; the lower laser power was used to avoid activating the neurotensin-expressing neurons in the nearby lateral hypothalamus. For all optogenetic manipulation experiment, inter-trial interval was set randomly from a uniform distribution between 7 and 15 min. Each experimental session lasted for 3-5 h (10-20 trials), and each animal was tested for 6-10 sessions.

### Chemogenetic manipulation

Saline (0.9% NaCl) or CNO (1 mg/kg, dissolved in saline) was injected intraperitoneally (i.p.) into mice expressing hM3Dq-mCherry, hM4Di-mCherry, or mCherry alone in the SNr. Each recording session started at 15 min after injection and lasted for 5 hours. Each mouse was recorded for 6-8 sessions, and CNO was given randomly in half of the sessions and saline in the other half. Data were averaged across all sessions for statistical comparison. For chemogenetic activation experiments in the new cage, each mouse was recorded for 2 sessions. Half of them were injected with CNO first and the other half with saline first.

### *In vivo* recording in the thalamus

To record single unit activity from thalamic neurons, we used custom-built optrodes, consisting of 12–14 wires (Stablohm 675, California Wire Company) twisted into stereotrodes and attached to an optic fiber (0.2 mm diameter). The electrodes were plated to an impedance of ∼ 200 kΩ and inserted into a screw-driven microdrive. We implanted the optrode into the VM of thalamus and used the microdrive to adjust the depth from 3.7 mm to 4.2 mm. Five second constant light was applied every 60 s for 60-80 trials.

Spikes were sorted as described previously(Weber et al., 2015). In brief, three largest principal components for each spike waveform on each stereotrode were used for offline sorting. Single units were identified either manually using the software Klusters (http://neurosuite.sourceforge.net) or automatically using the software KlustaKwik (http://klustakwik.sourceforge.net). Units with a clear refractory period in the auto-correlogram and satisfying criteria for isolation distance (> 20) and L-ratio (< 0.15) were regarded as single units.

### Transition analysis

To quantify transition probabilities between brain states, we discretized time into 15 s bins and aligned all laser stimulation trials from all mice by the onset of laser stimulation as time 0. To determine the transition probability from state X to Y for time bin i, Pi(Y|X), we first determined the number of trials (n) in which the animal was in brain state X during the preceding time bin i–1. Next, we identified the subset of these trials (m) in which the animal transitioned into state Y in the current time bin i. The transition probability Pi(Y|X) was computed as m/n. In Supplementary Fig.2a-b, each bar represents the transition probability averaged across four consecutive 15 s bins. To compute the baseline transition probabilities, we averaged across the 2 min period before laser stimulation.

To quantify transition probabilities between motor behaviors, we discretized time into 5 s bins. We calculated the effects of optogenetic manipulation by comparing the average transition probabilities during laser stimulation with that during the baseline period. In Supplementary Fig.3a-c each bar represents the transition probability averaged across twelve consecutive 5-s bins.

### Statistics

No statistical methods were used to pre-determine sample sizes, but our sample sizes are similar to those in previously published studies. No data point was excluded. For optogenetic and chemogenetic experiments, mice were randomly assigned to control (injected with AAV-DIO-eYFP or AAV-DIO-mCherry) and experimental groups. No randomization was used for rabies-mediated transsynaptic tracing. For mouse behavioral scoring, the scorers were blind to the animal identity and outcome assessment.

Bootstrap was used for testing the effects of laser stimulation on brain states and behavioral states and transition probabilities in all optogenetic experiments. Two-way ANOVA with Bonferroni correction was used for comparisons of brain state between different groups in chemogenetic experiments. Two-sided Wilcoxon signed-rank test was used for comparison of nest quality between different groups, and for comparison of firing rate of thalamic neurons during and before light stimulation. Two-sample Kolmogorov-Smirnov test was used to test the cumulative time for each behavior in new cage during the first hour. For all tests, we ensured that the variances of the data were similar between the groups under comparison.

The 95% confidence intervals (CIs) for percentage of time in each brain state or transition probability between each pair of brain state were calculated using a bootstrap procedure. For an experimental group of *n* mice, we calculated the CI as follows: we repeatedly resampled the data by randomly drawing for each mouse *p* trials (random sampling with replacement, *p* is the number of trials tested for that mouse). For each of the *m* iterations, we recalculated the mean across the *n* mice. The lower and upper CIs were then extracted from the distribution of the *m* resampled mean values. To test whether a given brain state or transition probability was significantly modulated by laser stimulation, we calculated for each bootstrap iteration the difference between the value during laser stimulation and that during the baseline period without laser stimulation. From the resulting distribution of difference values, we then calculated the *P-*value to assess whether laser stimulation significantly modulated brain states or transitions between brain states.

## Acknowledgments

We thank F. Weber and M. Xu for kindly sharing Matlab programs for data analysis, W. Chang for providing rabies virus for tracing experiments; J. Ding for helpful comments and discussions. This work was supported by Howard Hughes Medical Institute.

## Author Contributions

D.L. and Y.D. conceived and designed the experiments. D.L. performed most of experiments. C.M. performed a subset of rabies tracing experiments. W.Z. scored mouse motor behavior. C.M. and Y.Y. performed FISH experiments. D.L. and Y.D. wrote the manuscript.

## Materials & Correspondence

All correspondence and material requests should be made to Y.D.

## Supplementary Figures

**Supplementary Figure 1.**
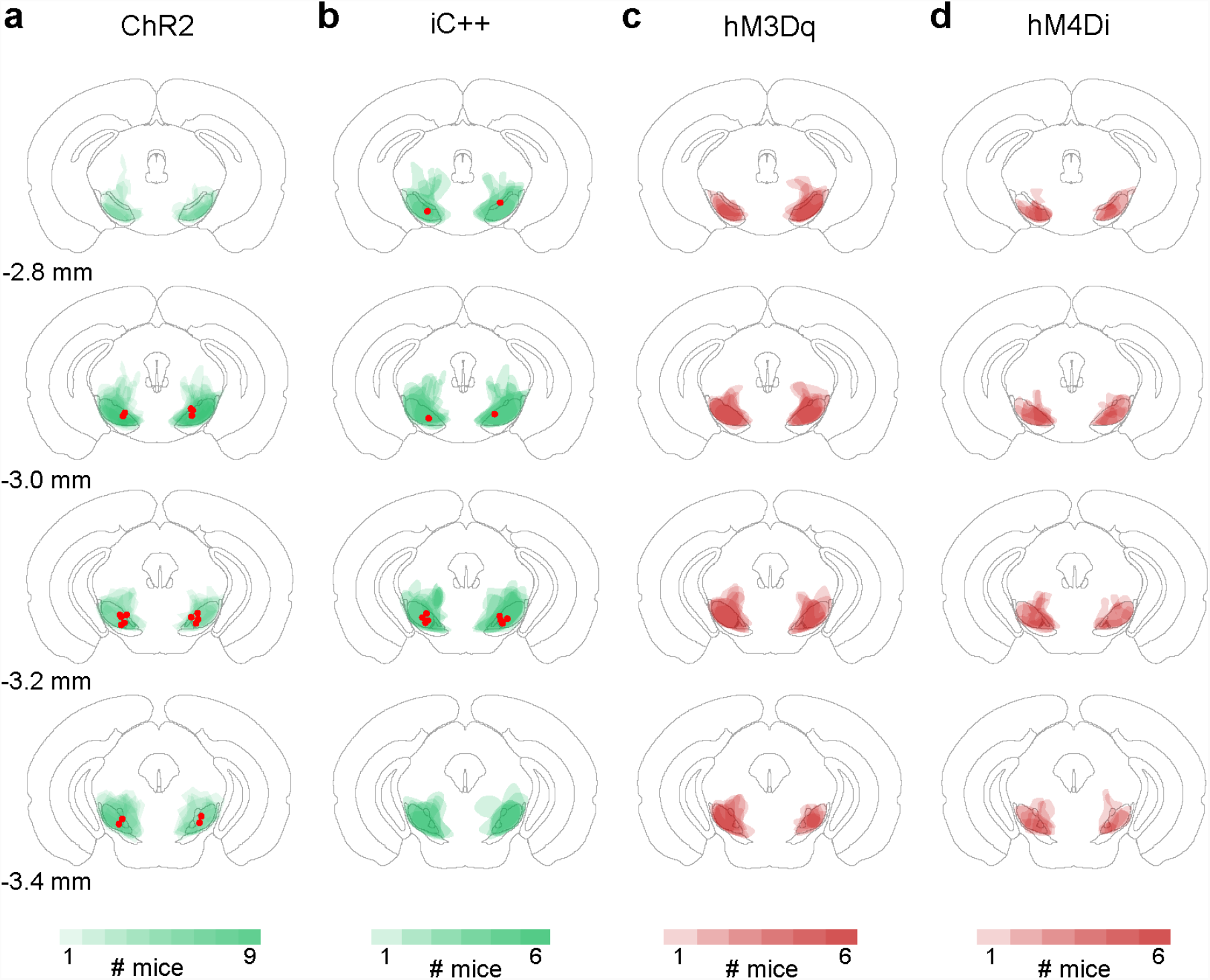
Locations of SNr GABAergic neurons targeted for optogenetic and chemogenetic manipulations. **a,** Summary of ChR2-eYFP expression in *Slc32a1*^*Cre*^ mice. For each mouse (n = 9), we determined the spread of ChR2-eYFP expression in 4 consecutive brain sections (from -2.8 mm to -3.4 mm along the rostrocaudal axis, where most of the virus expression was observed). The green color code indicates in how many mice the virus was expressed at the corresponding location. Red dots indicate the locations of optic fiber tips. Brain outline adapted from Allen Mouse Brain Atlas (available from: http://brain-map.org/api/index.html). **b-d,** Similar to **a,** for iC++-eYFP (b), hM3Dq-mCherry (c) and hM4Di-mCherry (d).

**Supplementary Figure 2.**
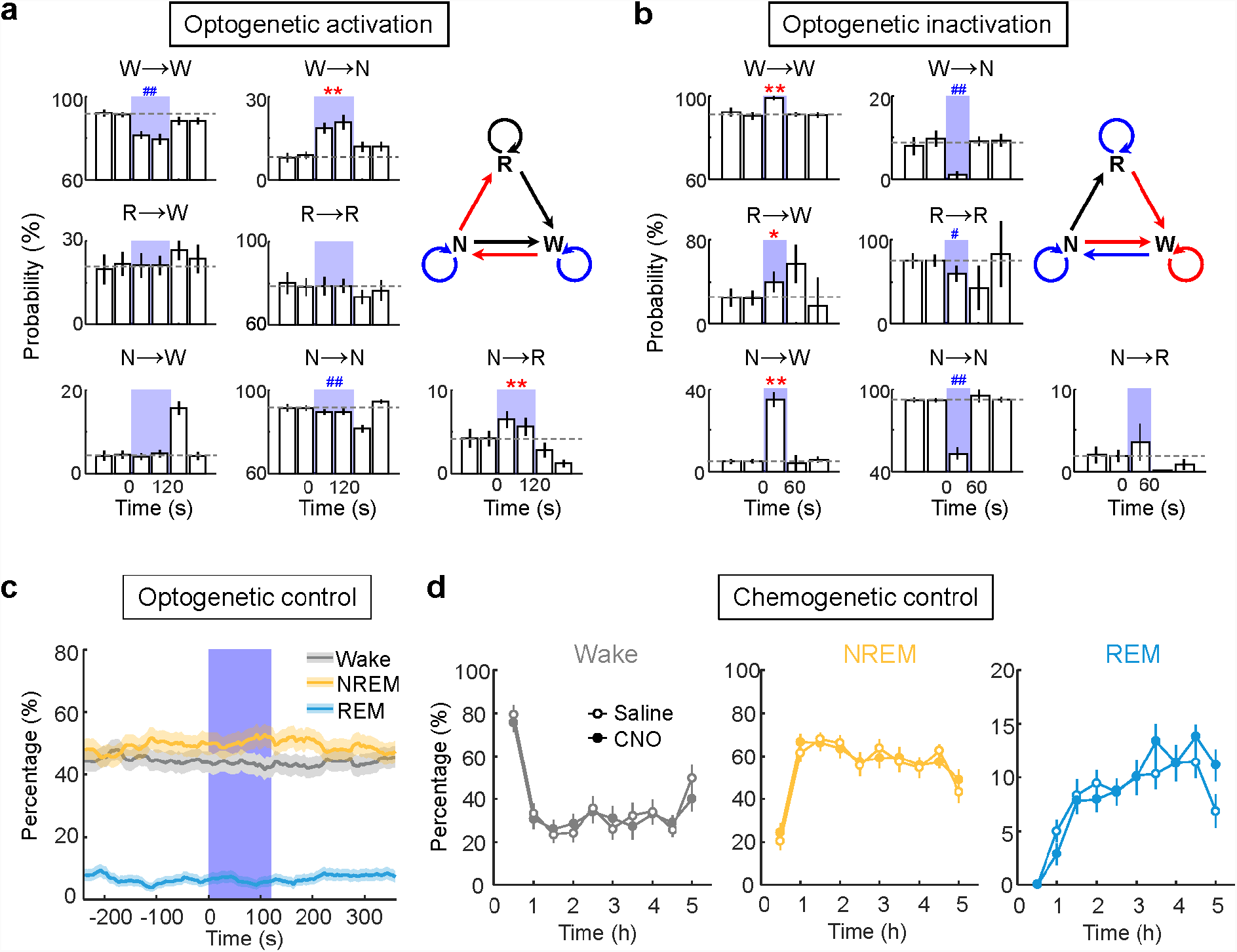
Effects of SNr activation and inactivation on brain state transition probability, and control experiment. **a,** Effect of optogenetic activation of SNr GABAergic neurons on transition probability between each pair of brain states. Each bar represents transition probability within each 15 s period (averaged across 4 consecutive 15 s bins within each 60 s). Error bar, 95% CI. Dashed line, baseline transition probability. N, NREM; R, REM; W, wake. W R and R N transitions are omitted because they were rarely detected with or without laser stimulation. */# indicates significant increase/decrease in transition probability during laser stimulation compared to baseline (*/# *P* < 0.05, **/## *P* < 0.001, bootstrap). The diagram summarizes transitions that are significantly increased (red), decreased (blue) or unchanged (black) during laser stimulation. **b,** Similar to **a**, for optogenetic inactivation. **c**, Effect of laser stimulation (constant light, 120 s) in *Slc32a1*^*Cre*^-eYFP control mice. Shown is the percentage of time in NREM, REM, or wake state before, during, and after laser stimulation, averaged from 6 mice (*P* > 0.1 for each brain state). Shading, 95% CI. Blue stripe, laser stimulation period. **d,** Effect of CNO in control *Slc32a1*^*Cre*^ mice expressing mCherry only. Shown is the percentage of time in each brain state following CNO or saline injection. Horizontal axis, time after CNO/saline injection. Error bar, ±SEM (n = 6 mice, 6-8 recording sessions/mouse). There was no significant difference between CNO and saline injections (*P* > 0.9, 0.9 and 0.4 for wake, NREM and REM respectively for all data points; two-way ANOVA with Bonferroni correction).

**Supplementary Figure 3.**
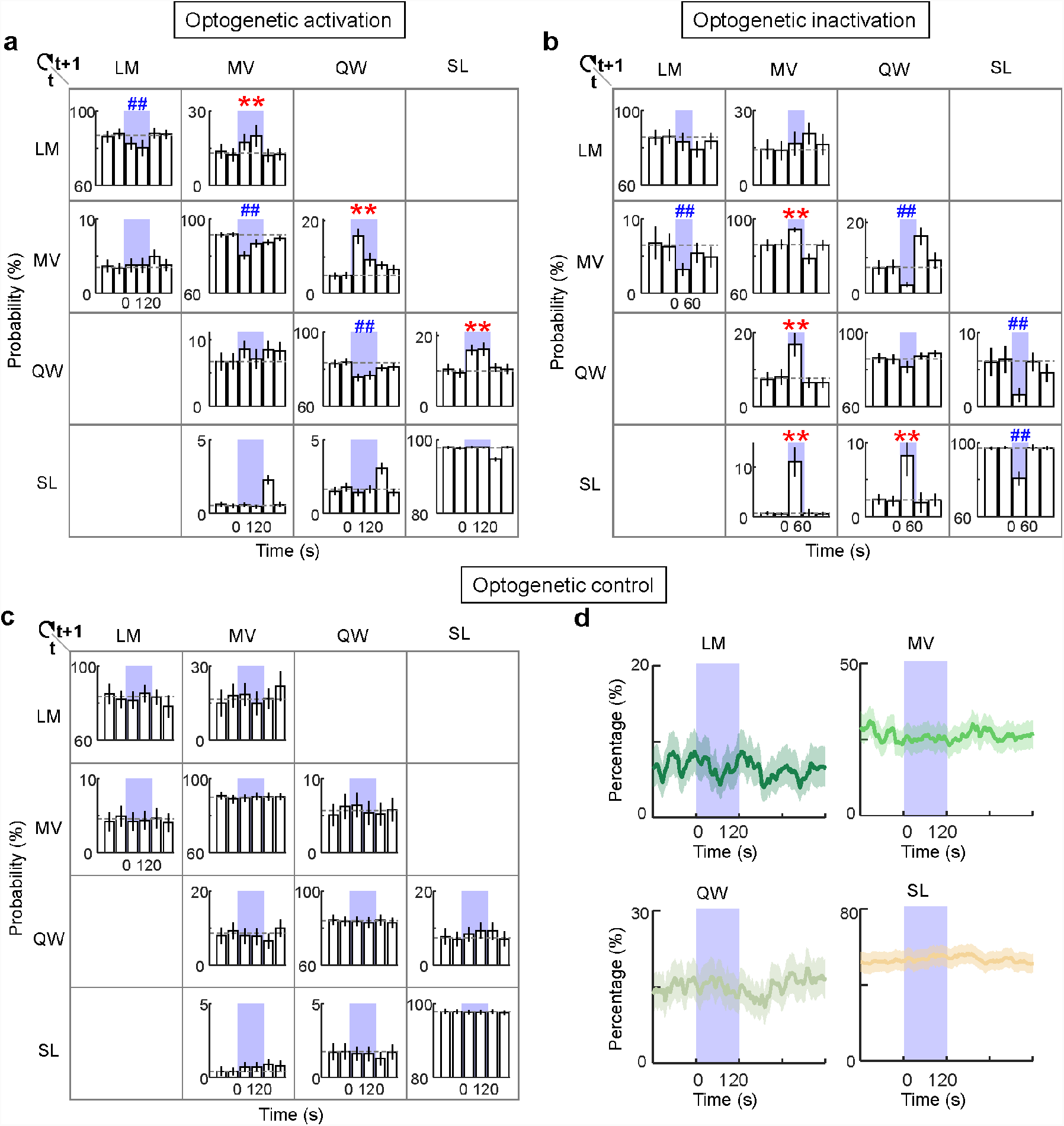
Effects of SNr activation and inactivation on probability of transition between behavioral states in home cage. **a,** Effect of optogenetic activation of SNr GABAergic neurons on transition probability between each pair of behavioral states. Each bar represents transition probability within each 5 s period (averaged across 12 consecutive 5 s bins within each 60 s). Error bar, 95% CI. Dashed line, baseline transition probability. */# indicates significant increase/decrease in transition probability during laser stimulation compared to baseline (*/# *P* < 0.05, **/## *P* < 0.001, bootstrap). **b & c,** Similar to **a**, for optogenetic inactivation in *Slc32a1*^*Cre*^-iC++ mice (b) and control experiment in *Slc32a1*^*Cre*^-eYFP mice (c). **d,** Effect of laser stimulation on the percentage of time in LM, MV, QW and SL states in *Slc32a1*^*Cre*^-eYFP control mice, averaged from 4 mice.

**Supplementary Figure 4.**
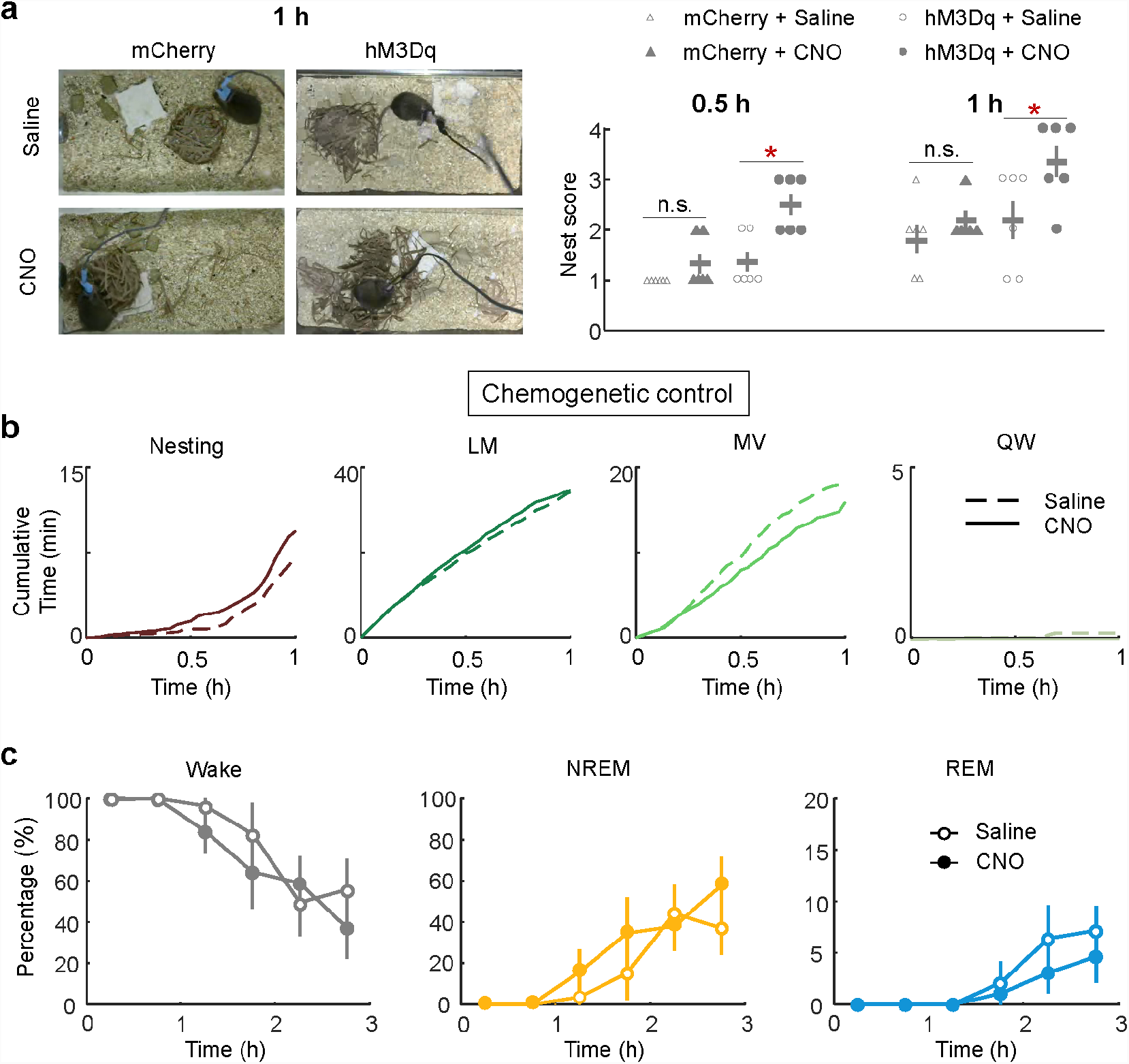
Effect of chemogenetic activation of SNr GABAergic neurons on nest quality in new cage, and control experiment. **a,** Effect of chemogenetic activation of SNr neurons on nest score (0: poor, 5: good). Left, example images of the nest 1 h after the mouse entered the new cage, for *Slc32a1*^*Cre*^-hM3Dq and *Slc32a1*^*Cre*^-mCherry mice after CNO or saline injection. Right, summary of nest scores at 0.5 h and 1 h for each group. Nest score was significantly higher for the hM3Dq group injected with CNO compared to saline (*P* < 0.05, Wilcoxon signed rank test), and compared to mCherry controls (*P* < 0.05, Wilcoxon ranksum test). No difference was found between CNO and saline for mCherry control mice. **b,** Effects of CNO on motor behaviors in *Slc32a1*^*Cre*^ mice expressing mCherry only. Shown is the cumulative time spent in each behavior after entering the new cage, averaged from 6 mice. There was no significant difference between CNO and saline injections (*P* > 0.1, two-sample Kolmogorov-Smirnov test). **c,** Effect of CNO on sleep and wakefulness in *Slc32a1*^*Cre*^ mice expressing mCherry only in the new cage. There was no significant difference between CNO and saline injections (*P* > 0.4, 0.1 and 0.1 for wake, NREM and REM, two-way ANOVA with Bonferroni correction).

**Supplementary Figure 5.**
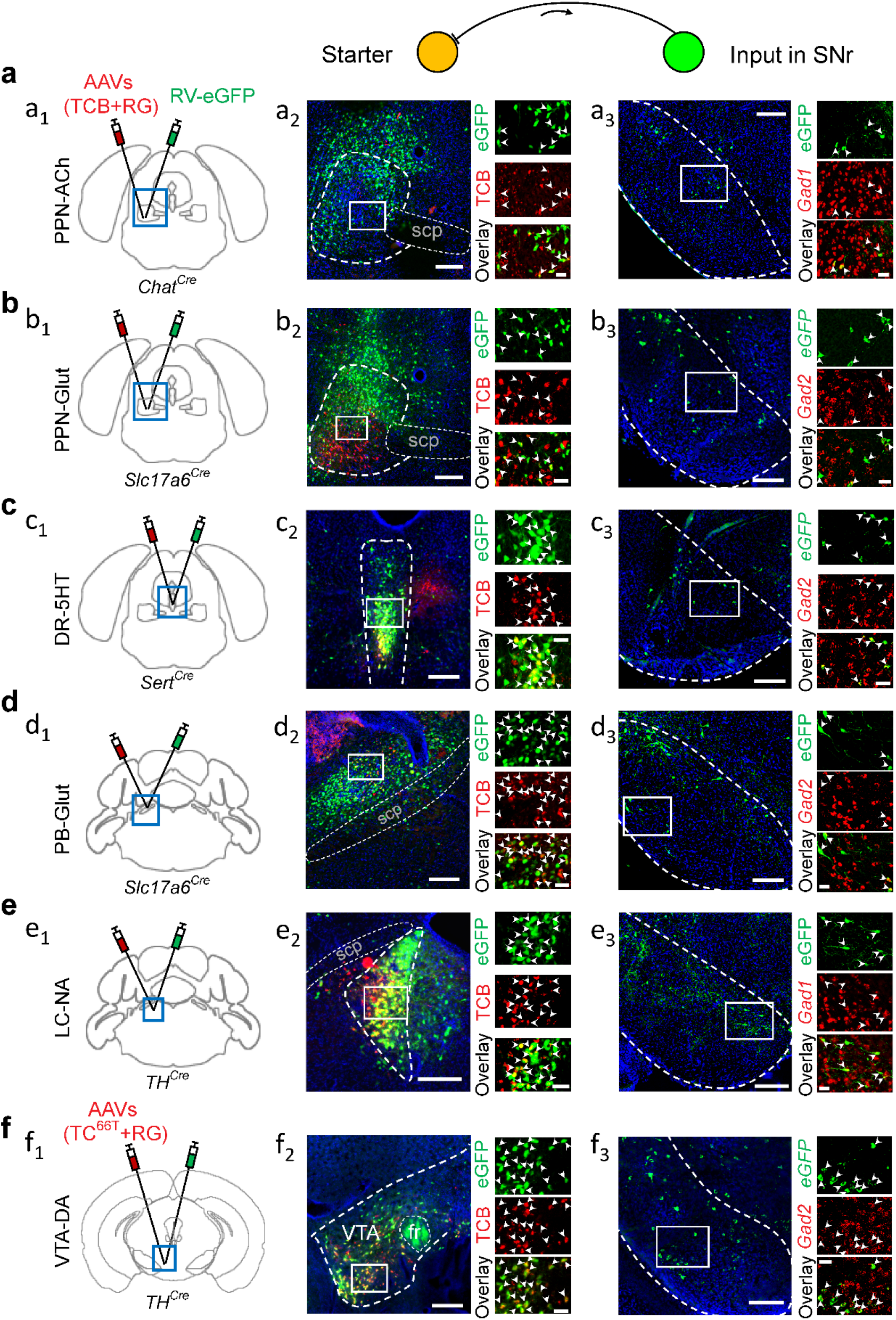
Innervation of multiple wake-promoting neuronal populations by SNr GABAergic neurons revealed by rabies-mediated circuit tracing. **a,** (**a_1_**), Schematic for rabies-mediated transsynaptic tracing from cholinergic (ACh) neurons in the pedunculopontine nucleus (PPN). (**a_2_**), Fluorescence image of PPN (box in coronal diagram in **a_1_**) together with enlarged view of the region in white box showing starter cells (expressing both eGFP and mCherry, indicated by arrowheads); scale bars, 200 μm and 25 μm. (**a_3_**), Rabies-labelled presynaptic neurons in SNr. Scale bar, 200 μm. Right images, enlarged view of the region in the white box, containing eGFP-labelled neurons expressing *Gad1*/*Gad2* (arrowheads); scale bar, 25 μm. **b-f,** Similar to **a**, for tracing from glutamatergic neurons in the PPN (**b**), serotonin neurons in the dorsal raphe (DR, **c**), glutamatergic neurons in the parabrachial nucleus (PB, **d**) and noradrenergic neurons in the locus coeruleus (LC, **e**), and dopaminergic neurons in the ventral tegmental area (VTA, **f**).

**Supplementary Figure 6.**
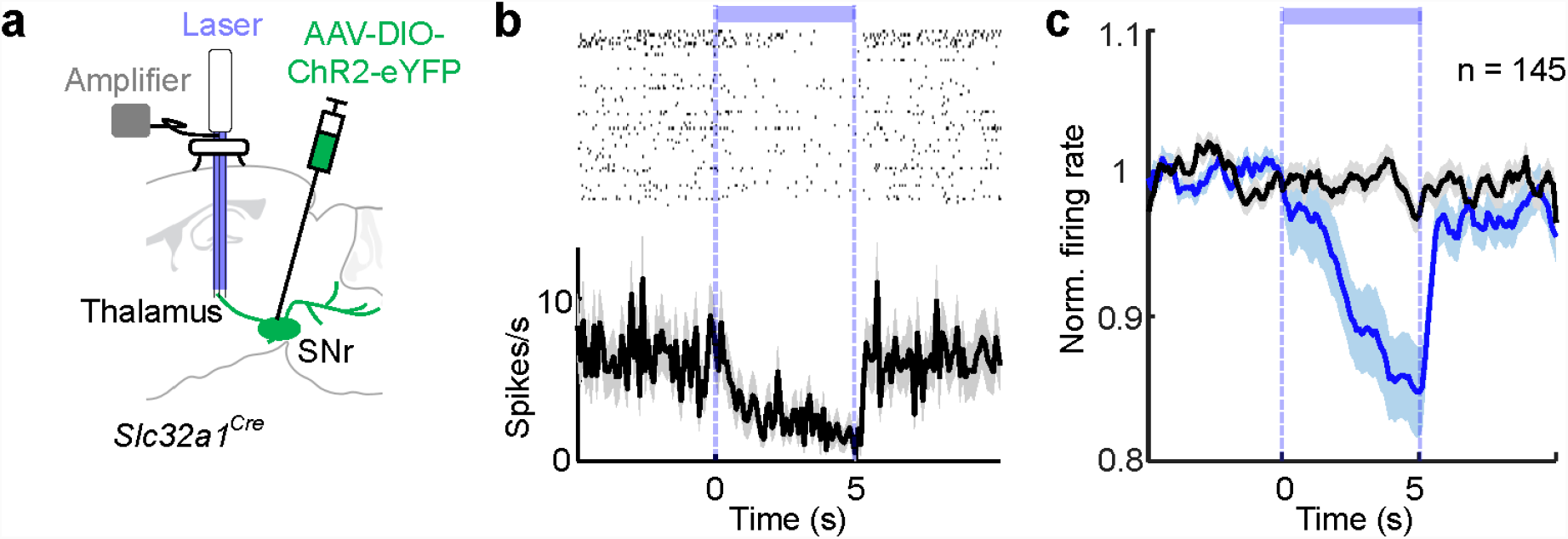
Activation of SNr axon terminals in the thalamus decreases thalamic firing rate. **a,** Schematic of optrode recording in the thalamus. **b,** Firing rate of an example unit in the thalamus before, during and after 5 s of laser stimulation of SNr GABAergic terminals in the thalamus. **c,** Average firing rate of 145 units from 4 mice (Blue, laser on; black, average from randomly selected laser off period). The firing rate of each unit was normalized by its baseline firing rate (before laser onset) before averaging across units. *P* < 0.0001 during laser-on, Wilcoxon signed rank test. Shading: SEM.

**Supplementary Figure 7.**
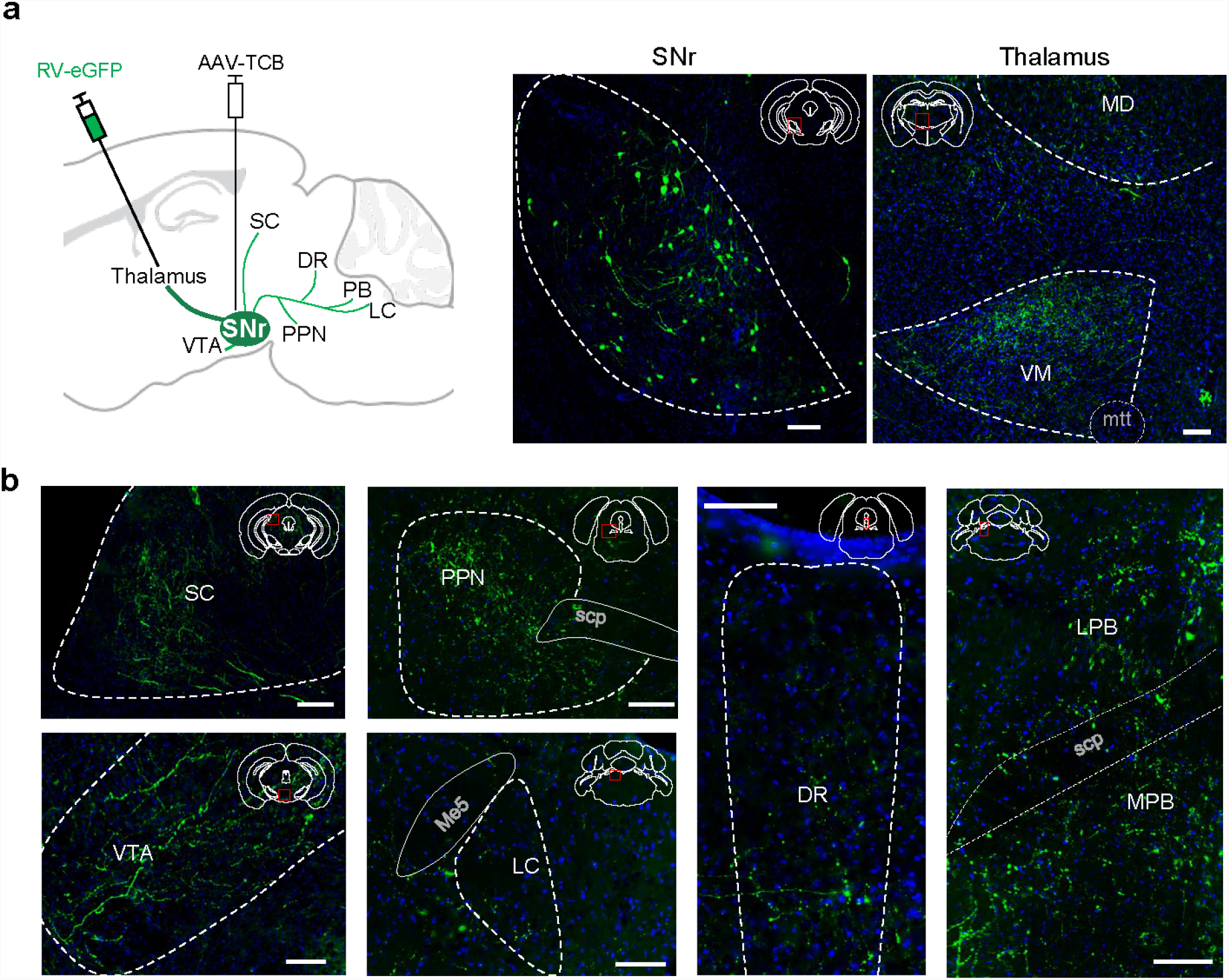
Axon collaterals of SNr GABAergic neurons that project to the thalamus. **a,** Left, schematic of viral strategy for labeling axon collaterals of thalamus-projecting SNr GABAergic neurons. AAV-DIO-TCB was injected into the SNr of *Slc32a1*^*Cre*^ mice and RV-eGFP injected into the thalamus. Middle, RV-eGFP-labelled neurons in the SNr. Right, axons in the thalamus. Scale bar, 100 μm; green, eGFP. **b,** Fluorescence images showing axon collaterals in the superior colliculus (SC), PPN, VTA, LC, DR, and PB. Scp, superior cerebellar peduncles; scale bar, 100 μm; green, eGFP.

**Supplementary Figure 8.**
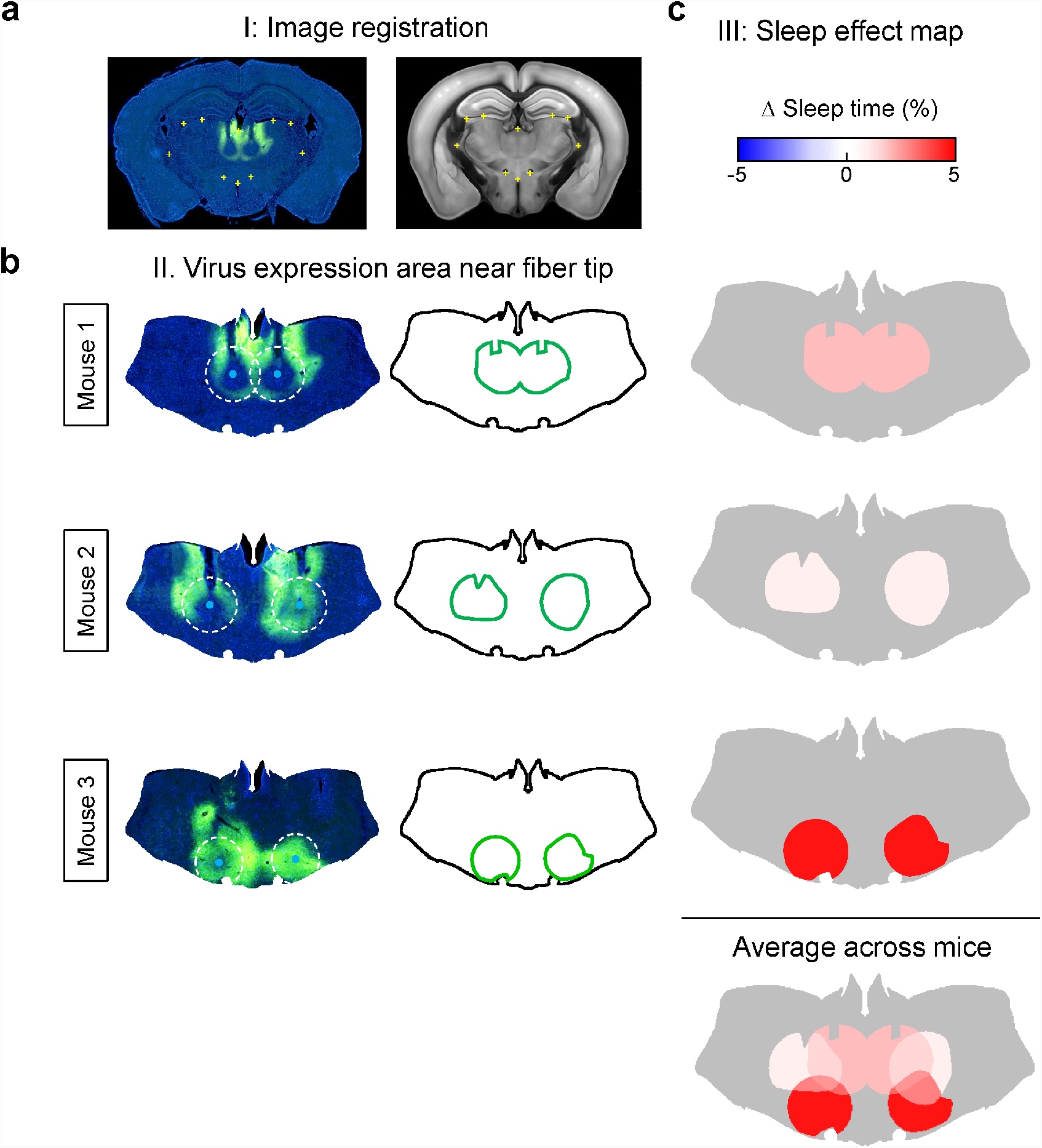
Procedure for analyzing the effect of optogenetic inactivation of thalamic neurons on brain states as a function of location. **a,** Image registration. Left, fluorescence image of a coronal section showing the expression of iC++-eYFP in a *Slc17a6*^*Cre*^ mouse. This image was aligned to the template from the Allen Mouse Brain Atlas (available from: http://brain-map.org/api/index.html) using multiple anatomical landmarks (yellow crosses) around or within the thalamus. **b,** Determination of virus expression area near the fiber tip. Left, images from three example mice at the same AP location. Dashed circle, area within 500 μm from fiber tip. Since laser stimulation normally causes eYFP photobleaching, we sacrificed the mice immediately after the last optogenetic testing session and used the center of the photobleached area together with the optic fiber track to determine the location of the fiber tip. After mapping the viral expression area based on fluorescence, the photobleached area was manually filled. Signals outside the sphere (500 μm in radius) surrounding the fiber tip were excluded. Right, green outline indicates area with viral expression near the fiber tip. **c,** Generation of sleep effect map. For each mouse, the effect of optogenetic inactivation on sleep (NREM and REM) was color-coded in the area with iC++-YFP expression near the fiber tip (green outline in **b**). The sleep effect for each location was calculated by averaging across all mice in which iC++-YFP was detected at that location (see bottom image, averaged over the three example mice). Gray: untested area in thalamus.

**Supplementary Figure 9.**
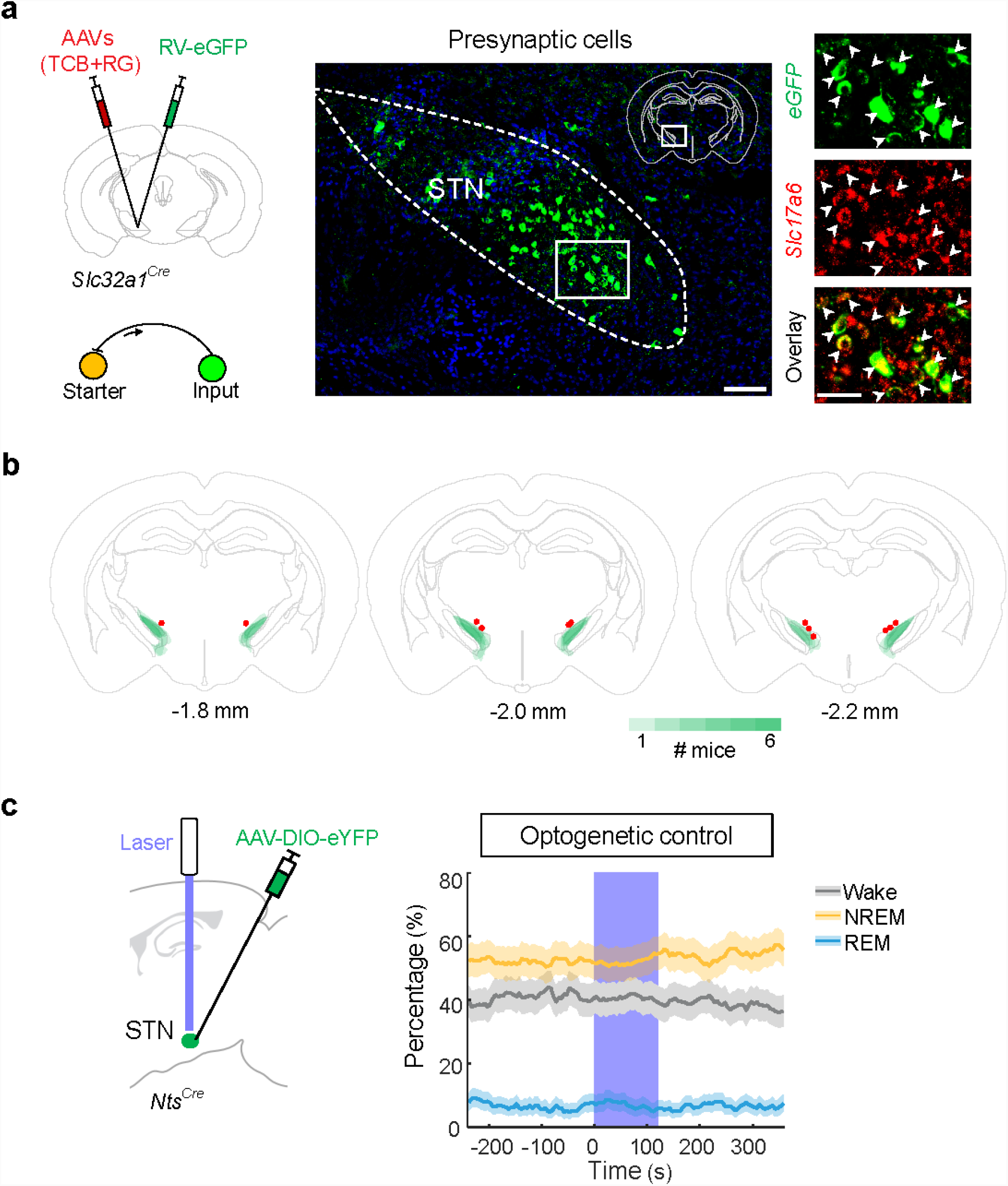
Innervation of SNr by STN glutamatergic neurons, locations of STN neurons targeted for optogenetic activation and control for STN optogenetic experiment. **a,** Left, schematic for rabies-mediated transsynaptic tracing from SNr GABAergic neurons. Middle, fluorescence image showing rabies-labelled presynaptic neurons in the STN (box in coronal diagram). Scale bar, 100 μm. Right, enlarged view of the region in the white box, containing eGFP-labelled neurons expressing *Slc17a6* (arrowheads); scale bar, 50 μm. **b,** Summary of ChR2-eYFP expression in the STN of *Nts*^*Cre*^ mice. For each mouse (n = 6), we determined the spread of ChR2-eYFP in 3 consecutive brain sections (from -1.8 mm to -2.2 mm along the rostrocaudal axis, where most of the virus expression was observed). The green color code indicates in how many mice the virus was expressed at the corresponding location. Red dots indicate the locations of optic fiber tips. **c,** Left, schematic of control experiment for STN activation. Right, effect of laser stimulation in NTS-eYFP control mice. Shown is the percentage of time in NREM, REM, or wake state before, during, and after laser stimulation (50 Hz, 120 s), averaged from 4 mice (*P* > 0.3 for all three brain states, bootstrap). Shading, 95% CI. Blue stripe, laser stimulation period.

